# SPHIRE-crYOLO: A fast and accurate fully automated particle picker for cryo-EM

**DOI:** 10.1101/356584

**Authors:** Thorsten Wagner, Felipe Merino, Markus Stabrin, Toshio Moriya, Claudia Antoni, Amir Apelbaum, Philine Hagel, Oleg Sitsel, Tobias Raisch, Daniel Prumbaum, Dennis Quentin, Daniel Roderer, Sebastian Tacke, Birte Siebolds, Evelyn Schubert, Tanvir R. Shaikh, Pascal Lill, Christos Gatsogiannis, Stefan Raunser

## Abstract

Selecting particles from digital micrographs is an essential step in single particle electron cryomicroscopy (cryo-EM). Since manual selection of complete datasets typically comprising many thousands of particles is a tedious and time-consuming process, many automatic particle pickers have been developed in the past few decades. However, non-ideal datasets pose a challenge to particle picking. Here, we present a novel automated particle picking software called crYOLO, which is based on the deep learning object detection system “You Only Look Once” (YOLO). After training the network with 500 – 2,500 particles per dataset, it automatically recognizes particles with high recall and precision reaching a speed of up to five micrographs per second. Importantly, we demonstrate a powerful general network trained on more than 40 datasets to select previously unseen datasets, thus paving the way for completely automated “on-the-fly” cryo-EM data pre-processing during data acquisition. CrYOLO is available as a standalone program under http://sphire.mpg.de/ and will be part of the image processing workflow in SPHIRE.

## Introduction

In recent years, single particle electron cryomicroscopy (cryo-EM) has become one of the most important and versatile methods for investigating the structure and function of biological macromolecules. In single particle cryo-EM, many images of identical but randomly oriented macromolecular particles are selected from raw cryo-EM micrographs, then digitally aligned, classified, averaged, and back-projected to obtain a three-dimensional structural model of the macromolecule.

Since more than 100,000 particles often have to be selected for a near-atomic cryo-EM structure, numerous automatic particle picking procedures, often based on heuristic approaches, have been recently developed^1–7^. In a popular selection approach, called template matching, cross-correlation of the micrographs is performed against pre-calculated templates of the particle of interest. However, this procedure is error-prone and the parameter optimization is often complicated. Although it works with good data, it often fails when dealing with non-ideal datasets where particles overlap, are compositionally and conformationally heterogeneous or the background of the micrographs is contaminated with crystalline ice. In these cases, these algorithms are not able to detect particles with high enough confidence, and consequently either particles are missed, or many false-positives are selected which need to be removed again afterwards. Often, the last resort is manual selection of particles, which is laborious and time-intensive.

To solve this problem, two particle selection programs^8,9^, DeepEM by Wang et al.^8^ and DeepPicker by Zhu et al.^9^ have been recently developed which employ deep convolutional neural networks (CNN). CNNs are extremely successful in processing data with a grid-like topology^10^. For images, this is undoubtedly the state-of-the-art method for pattern recognition and object detection. Similar to learning the weights and biases of specific neurons in a multi-layer perceptron^11^, a CNN learns the elements of convolutional kernels. A convolution is an operation which calculates the weighted sum of the local neighborhood at a specific position in an image. The weights are the elements of the kernel which extracts specific local features of an image, such as corners or edges. In convolutional neural networks, several layers of convolutions are stacked, where the output of one layer is the input of the next layer. This enables CNNs to learn hierarchies of features thereby allowing the learning of complex patterns.

Like common modern object detection systems, particle selection tools employ a specific classifier for an object and evaluate it at every position. Therefore they are trained with positive examples of cropped out particles and negative examples of cropped out regions of background or contamination. After the classifier is trained, these systems slide a window over the micrograph to crop out single local regions, pass it through a CNN and finally classify the extracted region. The confidence of classification is transferred into a map and the object positions are estimated by finding the local maxima in this map. Using this approach, it is possible to select particles on more challenging datasets. However, as it classifies many overlapping cut-out regions independently, it comes with a high computational cost. Moreover, since the classifier only sees the windowed region, it is not able to learn the larger context of a particle (e.g. to not pick regions near ice contamination).

In 2016, Joseph Redmon and colleagues introduced the ‘You Only Look Once’ (YOLO) framework^12^ as an alternative to the sliding window approach and reformulated the classification problem into a regression task, where the exact position of the particle is predicted by the network. In contrast to the sliding window approach, the YOLO framework requires only a single pass of the full image instead of multiple passes of cropped-out regions. Thus, the YOLO approach essentially divides the image into a grid and predicts for each grid cell whether it contains a center of a bounding box enclosing an object of interest. If this is the case, it applies regression to predict the relative position of the object center inside the cell as well as the width and height of the bounding box (Figure 1a). This procedure makes the detection pipeline rather simple and reduces the number of convolutions tremendously, which increases its speed while remaining accurate. During training, the YOLO approach only requires labeling positive examples, whereas the sliding window approach also requires labeling background and contaminants as negative examples. Moreover, since the network sees the complete image, it is able to learn the larger context and therefore provides an excellent framework to reliably detect single particles in electron micrographs (Figure 1b).

**Figure 1.**
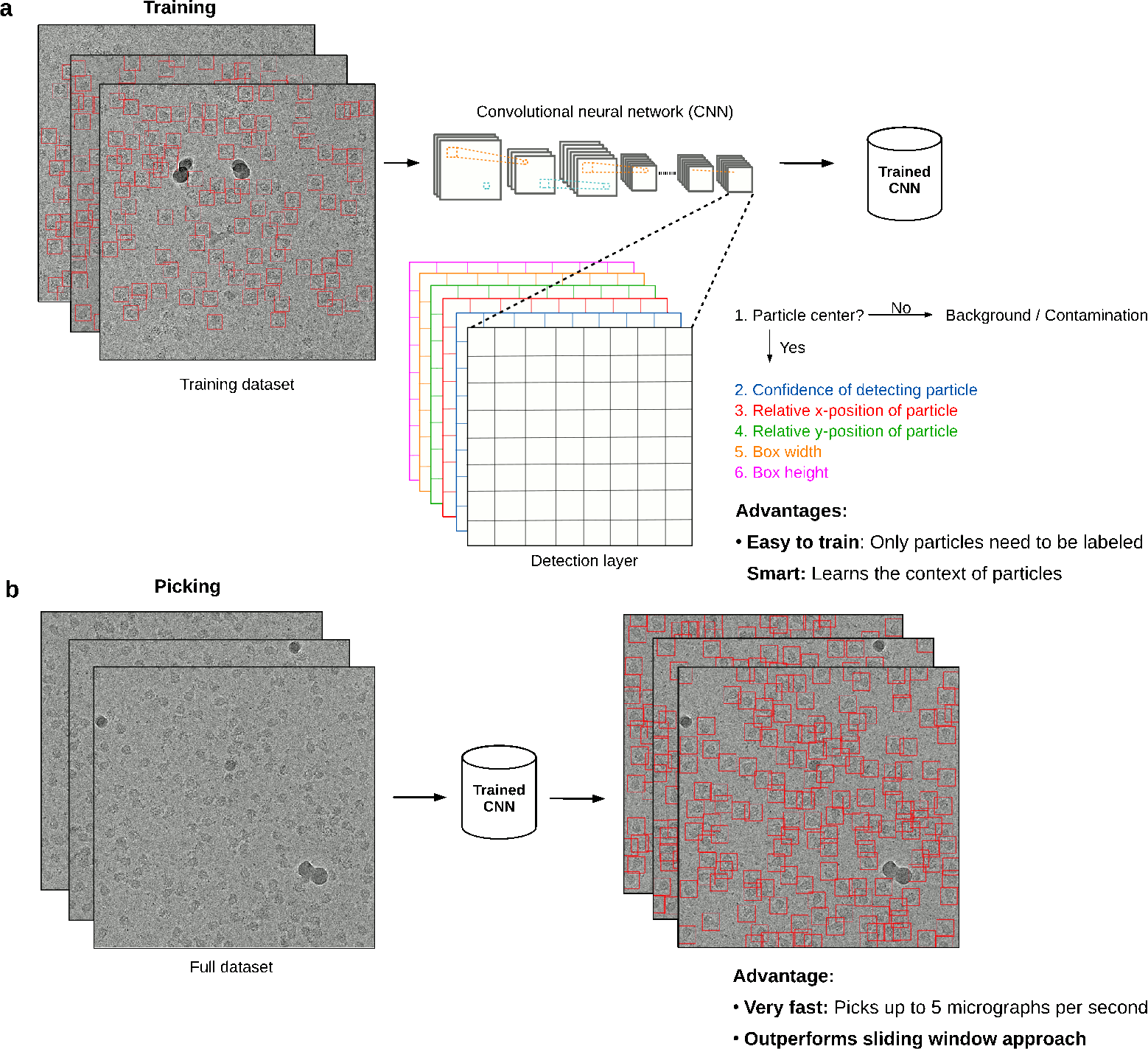
Training and picking in crYOLO. **a)** With YOLO approach, the complete micrograph is taken as the input for the CNN. When the image is passed through the network, the image is spatially downsampled to a small grid. Then, YOLO predicts for each grid cell if it contains the center of a particle bounding box. If this is the case, it estimates the relative position of the particle center inside the cell as well as the width and height of the bounding box. During training, the network only needs labeled particles. Furthermore, as the network sees the complete micrograph, it learns the context of the particle. **b)** The picking is very fast. It picks up to 5 micrographs per second and thus outperforms the sliding window approach.

Here, we present a novel single particle selection procedure crYOLO which utilizes the YOLO algorithm to select single particles in cryo-EM micrographs. We evaluate our procedure on simulated data, a common benchmark dataset and three recently published high resolution cryo-EM datasets. The results demonstrate that crYOLO is able to accurately and precisely select single particles in micrographs of varying quality with a speed of up to five micrographs per second on a single GPU. CrYOLO leads to a tremendous acceleration of particle selection, simplifying the training as no negative examples have to be labeled, and improving not only quality of the extracted particle images but also the final structure. Furthermore, we present a general model for crYOLO, which was trained on more than 40 datasets and is able to select particles of previously unseen macromolecular species, realizing automatic particle picking at expert level.

## Results

### Convolutional Neural Network

The program crYOLO builds upon a Python-based open source implementation^13^ of YOLO and uses the deep learning library Keras^14^. Beyond the basic implementation, we added patch-processing, multi-GPU support, parallel processing, preprocessing procedures, support for MRC micrographs, single channel data, RELION star files and EMAN1^15^ box files. CrYOLO includes a graphical tool to read and create box files for training data generation or visualization of the results in a user-friendly manner (Figure 2). For the readers who are interested in the details of YOLO architecture, please refer the online methods and the original paper^16^. Briefly, the YOLO network consists of 22 convolutional and five max-pooling layers. In addition, it contains a passthrough layer between layer 13 and 21 (similar to ResNets^17^) to exploit fine grain features and the network is followed by a (1×1) convolutional layer that is used as a detection layer. This architecture shows a similar performance to a 50-layer ResNet but with a much lower number of weights^16^. However, one limitation of the original YOLO network for particle picking is that it uses a relatively coarse grid for prediction. Under special circumstances, e.g. when particles are very small, this might lead to a lower performance due to the fact that each grid cell can only detect a single particle.

**Figure 2.**
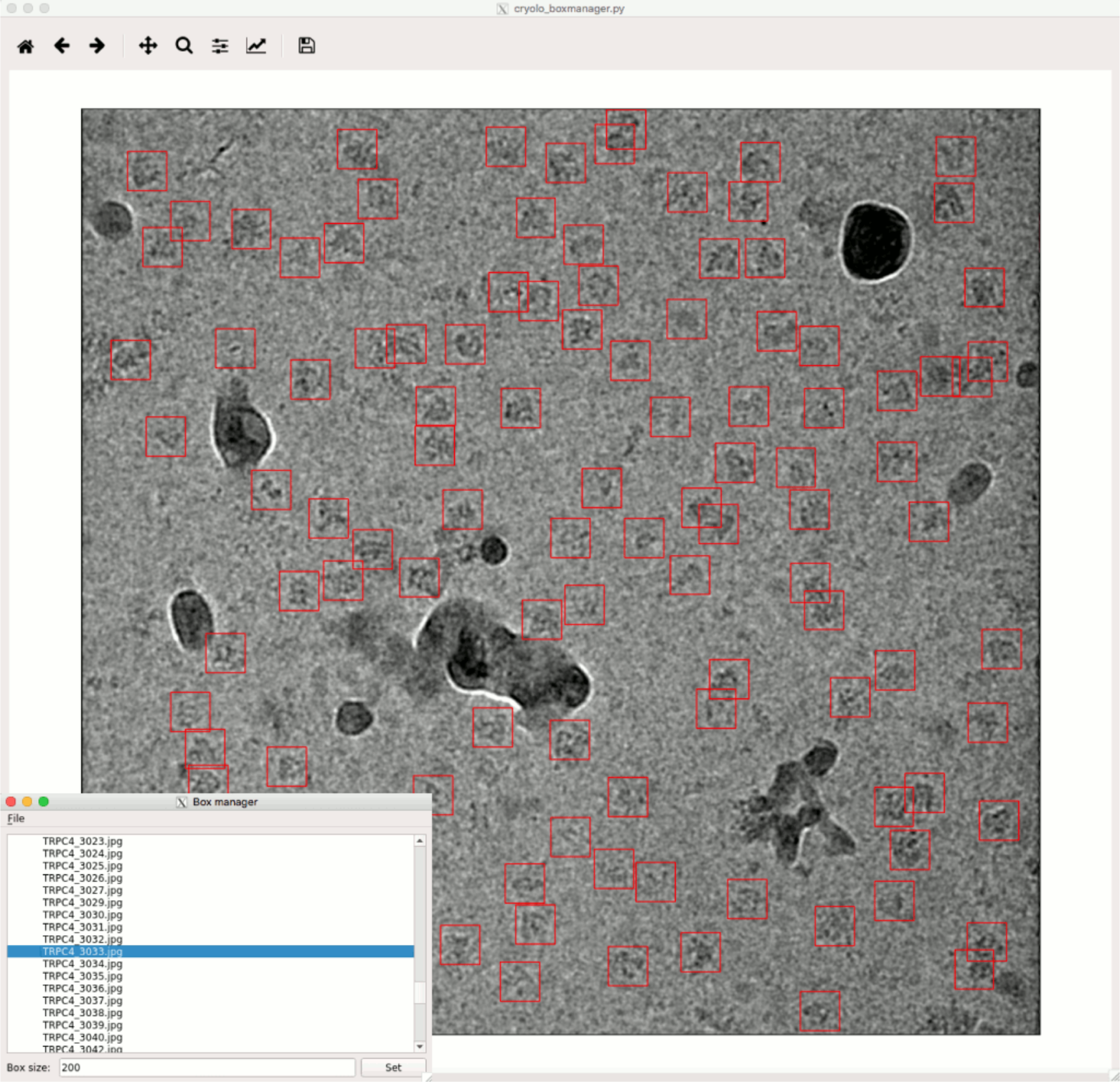
Graphical tool for creating training data and visualizing results. The tool can read images in MRC, TIFF and JPG format and box files in EMAN1 and STAR format. The example shown is a micrograph of TRPC4^21^ with many contaminants.

In order to tackle the problem of detecting very small particles, crYOLO divides the input image into a small number of overlapping patches (e.g. 2×2 or 3×3). Instead of the complete micrograph, each patch is then downscaled to the network input image size of 1024×1024. During prediction, all patches are classified in a single batch.

To reduce overfitting and improve the generalization of the trained network, each image is internally augmented before passed through the network during training. During augmentation a random selection of the following methods are used: flipping, blurring, adding noise and random contrast changes; which are described in the online methods.

### Test datasets

We used crYOLO to select three different cryo-EM datasets and analyzed the results. These include the TcdA1 toxin subunit from *Photorhabdus luminescens*^18^ (EMPIAR-10089, EMDB-3645), the *Drosophila* transient receptor channel NOMPC^19^ (EMPIAR-10093, EMDB-8702) and the human peroxiredoxin-3 (Prx3)^20^ (EMPIAR-10050, EMDB-3233). All datasets were previously shown to produce high-resolution structures. To validate the performance of our program we also tested it on simulated data of the canonical TRPC4 ion channel^21^ (EMD-4339) and on a published dataset of the keyhole limpet hemocyanin (KLH)^22^.

TcdA1 assembles in the soluble prepore state in a large bell-like shaped pentamer with a molecular weight of 1.4 MDa^18^. TcdA1 has been our test specimen to develop the software package SPHIRE^23^. We previously used a dataset of 7,475 particles of TcdA1 to obtain a reconstruction at a resolution of 3.5 Å^23^. Due to the large molecular weight of the specimen and its characteristic shape, particles are clearly discernible in the micrographs, although a carbon support film was used. Therefore, the selection of the particles is straightforward in this case. We chose this particular test dataset because of its small size, since the quality and number of selected particles will likely have an influence on the quality of the final reconstruction.

In the case of NOMPC, which was reconstituted in nanodiscs^19^, the overall shape of the particle, sample concentration, and limited contrast make it difficult to accurately select the particles, despite having a molecular weight of 754 kDa. Furthermore, the density of the nanodisc is more pronounced than the density of the extramembrane domain of the protein, thus the center of mass is not located at the center of particle but shifted towards the nanodisc density. We chose this dataset as it is a challenge for the selection program to accurately detect the center of the particles and to avoid selection of empty nanodiscs.

Prx3 has a molecular weight of 257 kDa and has a characteristic ring-like shape. The dataset is one of the first near-atomic resolution datasets recorded using a Volta phase plate^20^ (VPP). The VPP introduces an additional phase shift in the unscattered beam. This increases the phase contrast, providing a significant boost of the signal-to-noise ratio in the low-frequency range. This makes the structural analysis of low molecular weight complexes at high resolution possible and has revolutionized the cryo-EM field^24^. The VPP, however, enhances not only the contrast of the particles of interest, but also the contrast of all weak phase objects, including smaller particles (impurities, contamination, dissociated particle components) that would otherwise not be easily visible in conventional cryo-EM. This poses new challenges with regard to particle selection, especially for automated particle picking procedures that cross-correlate raw EM images with templates.

To obtain a simulated dataset, we produced sets of 20 micrographs, each with 250 randomly oriented TRPC4 particles, but with a different noise level *k*. To generate realistic noise, we followed a similar procedure as described in Baxter et al^25^. To simulate structural noise, we first added zero mean Gaussian noise with a standard deviation of 88\**k*. Then we applied a CTF with a fixed defocus of 2.5 micrometers, voltage of 300 kV, amplitude contrast of 0.1 and a B-factor of 200. Finally, to simulate shot and digitalization noise, we added mean free Gaussian noise again with a standard deviation of 51\**k*. With this procedure we produced 5 sets with *k* = 1, 4, 6, 8 and 10. The corresponding SNRs, defined by signal power divided by noise power, are 1.25, 0.093, 0.041, 0.023, 0.014 respectively.

A common benchmark dataset for particle picking is a published set of cryo-EM images of KLH^22^. Besides KLH particles, the dataset contains mitochondrial antiviral signaling (MAVS) filaments, stacked KLH particles and broken particles. Ideally, all of these contaminants would be ignored by a particle picking procedure and only intact and single KLH particles selected. An additional advantage of the dataset is, that the micrographs were recorded as pairs with high and low defocus, which allows to evaluate the performance of a particle picker for each defocus separately.

### Training and application of crYOLO

To train crYOLO, we manually selected particles for initial training datasets. Depending on the density of particles, the heterogeneity of the background, and the variation of defocus, more or fewer micrographs are needed. For the TcdA1, NOMPC and Prx3 datasets, we found that 500 to 2,500 particles from at least five micrographs were sufficient to properly train the networks for the three datasets. It should be emphasized that the picking of negative examples (including background, carbon edges, ice contamination and “bad particles”) is not required at all in crYOLO, as essentially all other positions are considered to be negative examples. It is sufficient that these contaminants are present in the training images with the labeled particles. Ideally, each micrograph should be picked to completion. However, since the contrast in cryo-EM micrographs is often low, a user is typically not able to select all particles for training and often misses some of them, referred to as false negatives. Including false negatives carries a lower penalty than missing true positives during training (see online methods). This enables crYOLO to converge during training, even if not all particles in a micrograph are picked.

To assess the performance of crYOLO we calculated precision and recall scores^26^. The scores were calculated on twenty percent of the micrographs that were used for manual selection, but not for training. The recall score measures the ability of the classifier to detect positive examples and the precision score specifies the strength of the classifier to not label a true negative as a true positive. Both scores are summarized with the integral of the precision-recall curves, the so-called area under the curve (AUC), which is an aggregate measure for the quality of detection. The larger the AUC value, the better the performance; 1.0 is the maximum at perfect performance.

In order to quantify how well the particles are centered, we calculated the mean intersection over union (IOU) value of manually selected particles versus the automatically picked boxes using crYOLO. The IOU is defined as a ratio of the intersecting area of two bounding boxes and the area of their union and is a common measure for the localization accuracy. Picked particles with an IOU higher than 0.6 were classified as true positive. The mean IOU for TcdA1, Prx3 and NOMPC was 0.86 ± 0.009 and 0.80 ± 0.004, 0.85 ± 0.007 respectively. This indicates a high localization accuracy of crYOLO.

To further assess the quality of particles picked by crYOLO, we additionally calculated 2-D classes using the iterative stable alignment and clustering approach^27^ (ISAC) as implemented in the SPHIRE software package^23^. 2-D clustering using ISAC relies on the concepts of stability and reproducibility and is capable of extracting validated and homogeneous subsets of images that can be reliably processed in steps further downstream in the workflow. The number of particles in these subsets indirectly reflects the performance of the particle picker. Moreover, we calculated 3-D reconstructions and compared it to the published reconstruction.

For TcdA1, crYOLO was trained on 10 micrographs with 1100 particles and selected 10,854 particles from 98 micrographs. This is ~29% more particles than previously identified by the Gauss-Boxer in EMAN2^15^ (Figure 3a-c). Furthermore, the number of ‘good’ particles after 2-D classification is higher (Figure 3d,e), indicating that crYOLO is able to identify more true positive particles. This results in a slightly improved resolution of the 3-D reconstruction (Figure 3f,g).

**Figure 3.**
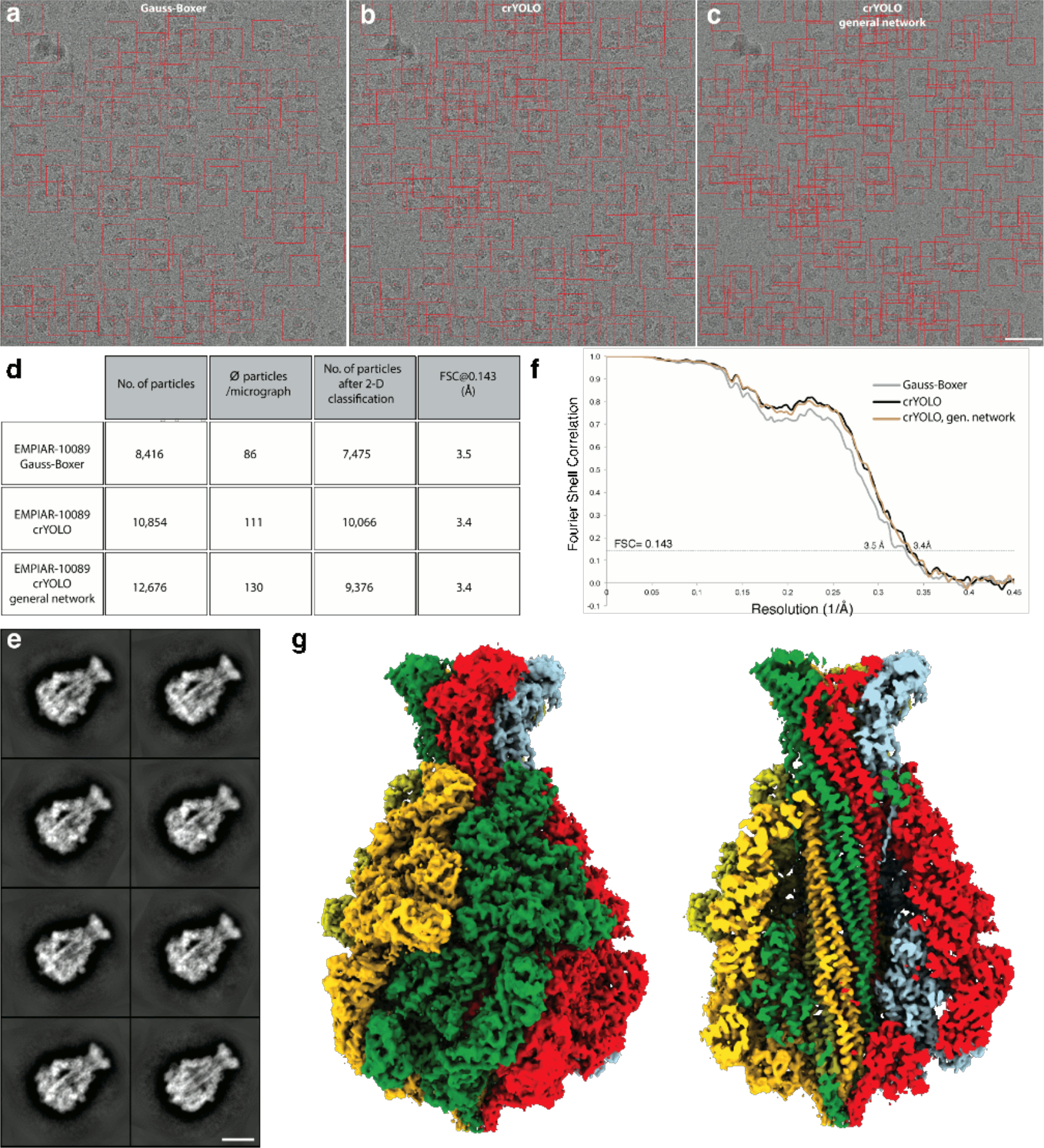
Selection of TcdA1 particles and structural analysis. **a-c**) Representative digital micrograph (micrograph number 0169) taken from the EMPIAR-10089 dataset. Red boxes indicate the particles selected by (**a**) Gauss-Boxer, (**b**) crYOLO or (**c)** the generalized crYOLO network. Scale bar, 50 nm. **d**) Summary of particle selection and structural analysis of the three datasets. All datasets were processed using the same workflow in SPHIRE. **e)** Representative reference-free 2-D class averages of TcdA1 obtained using the ISAC and Beautifier tools (SPHIRE) from particles picked using crYOLO. Scale bar, 10 nm. **f**) Fourier shell correlation (FSC) curves of the 3-D reconstructions calculated from the particles selected in crYOLO and Gauss-Boxer. The FSC 0.143 between the independently refined and masked half-maps indicates resolutions of ~3.4 and ~3.5 Å, respectively. **g)** The final density map of TcdA1 obtained from particles picked by crYOLO is shown from the side, and is colored by subunit. The reconstruction using particles from the generalized crYOLO network is indistinguishable.

For NOMPC, the original authors picked initially 337,716 particles from 1,873 micrographs using RELION^5^ (Figure 4b,c) and obtained a 3-D reconstruction at a resolution of 3.55 Å from 175,314 particles after extensive cleaning of the dataset by 2-D and 3-D classification using RELION^28^. CrYOLO was trained on 9 micrographs with 585 particles and selected only 226,289 particles (> 30% fewer) (Figure 4a,c). After 2-D classification and the removal of bad classes (Figure 4c,d), 171,917 particles were used for the 3-D refinement yielding a slightly improved cryo-EM structure of NOMPC of 3.4 Å (Figure 4e-f). The reconstruction shows high resolution features even in the region of the ankyrin-repeats (Figure 4g). Although the number of initially selected particles differed tremendously from crYOLO, the number of particles used for the final reconstruction is very similar, indicating that crYOLO selects less false positive particles than the selection tool in RELION. In this case, this also reduced the steps and overall time of image processing, since 2-D classification was performed on a much lower starting number of particles and further cleaning of the dataset by a laborious 3-D classification was unnecessary.

**Figure 4.**
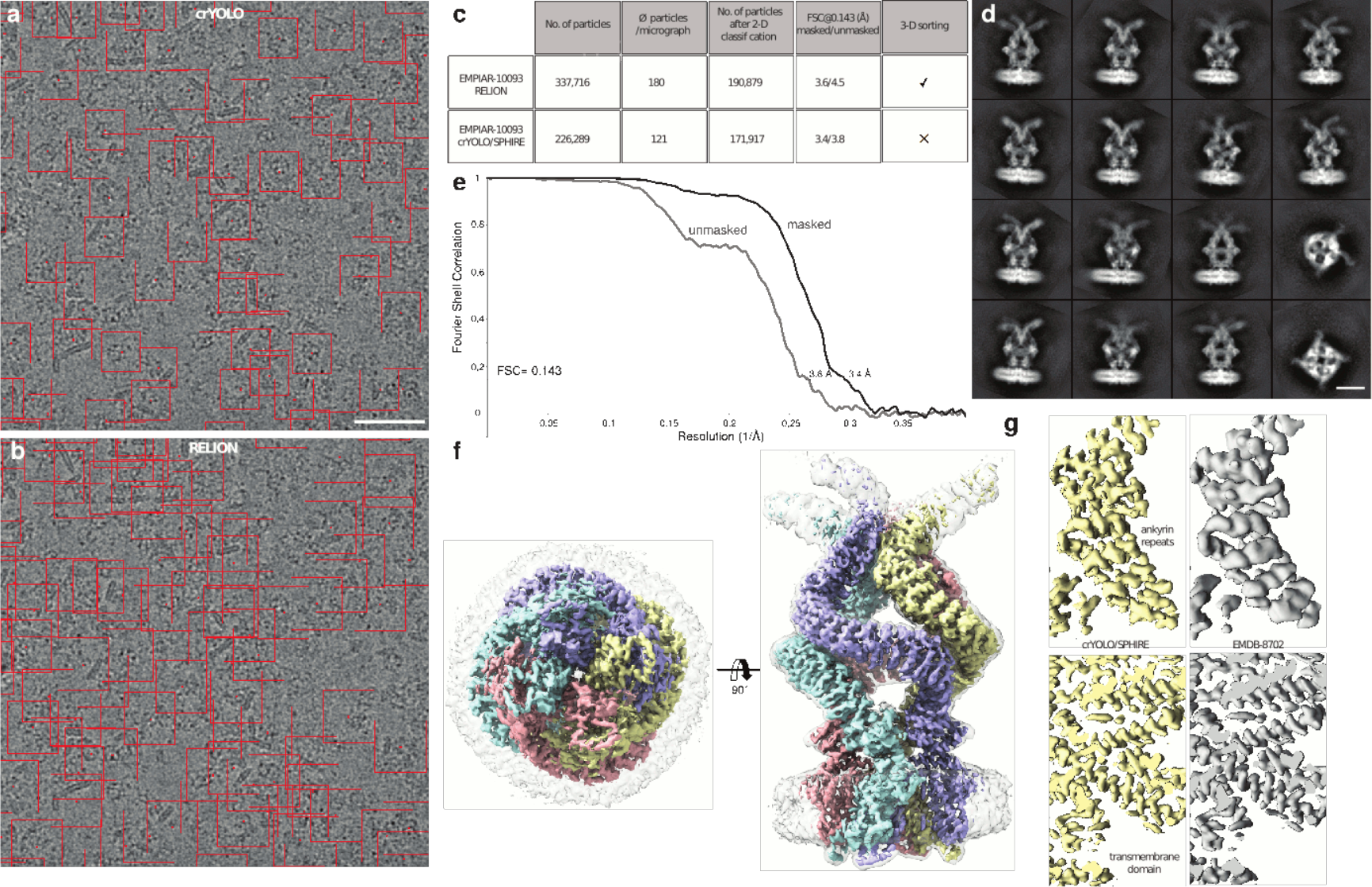
Selection of NOMPC particles and structural analysis. **a, b)** Representative micrograph (micrograph number 1,854) of the EMPIAR-10093 dataset. Particles picked by **(a)** crYOLO or **(b)** RELION, respectively, are highlighted by red boxes. Scale bar, 50 nm. **c)** Summary of particle selection and structural analysis using RELION and crYOLO/SPHIRE. **d)** Representative reference-free 2-D class averages obtained using the ISAC and Beautifier tools (SPHIRE) from particles selected by crYOLO. Scale bar, 10 nm. **e)** FSC curves **(f)** and final 3-D reconstruction of the NOMPC dataset obtained from particles picked using crYOLO and processed with SPHIRE. The 0.143 FSC between the masked and unmasked half-maps indicates resolutions of 3.4 and 3.8 Å, respectively. The 3-D reconstruction is shown from the top and side. To allow better visualization of the nanodisc density the unsharpened (gray, transparent) and sharpened map (colored by subunits) are overlaid. **g)** Comparison of the density map obtained by crYOLO/SPHIRE with the deposited NOMPC 3-D reconstruction EMDB-8702.

For Prx3, crYOLO was trained on 5 micrographs with 2500 particles and identified 354,581 particles from 802 digital micrographs, which is comparable to the number of particles picked in the original article using EMAN2^15^ (Figure 5a-c). We used two consecutive ISAC rounds to classify the particles in 2-D (Figure 5d). In the first one, we classified the whole dataset into large classes to identify the overall orientation of the particles. In the second one, we split them into top, side and tilted views, and ran independent classifications on each of them. This procedure is necessary to avoid centering errors in ISAC. After discarding bad classes and removing most of the preferential top view orientations, the dataset was reduced to 37,031 particles. 3-D refinement of this particle stack in SPHIRE yielded a map at a nominal resolution of 4.6 Å (Figure 5e, f). In comparison, the final stack in the original article was composed of only 8,562 particles, which were refined to a 4.4 Å resolution map in RELION. In contrast to our processing pipeline, the original authors used three rounds of 2-D and four rounds of 3-D classification in RELION^5^ in order to clean this dataset.

**Figure 5.**
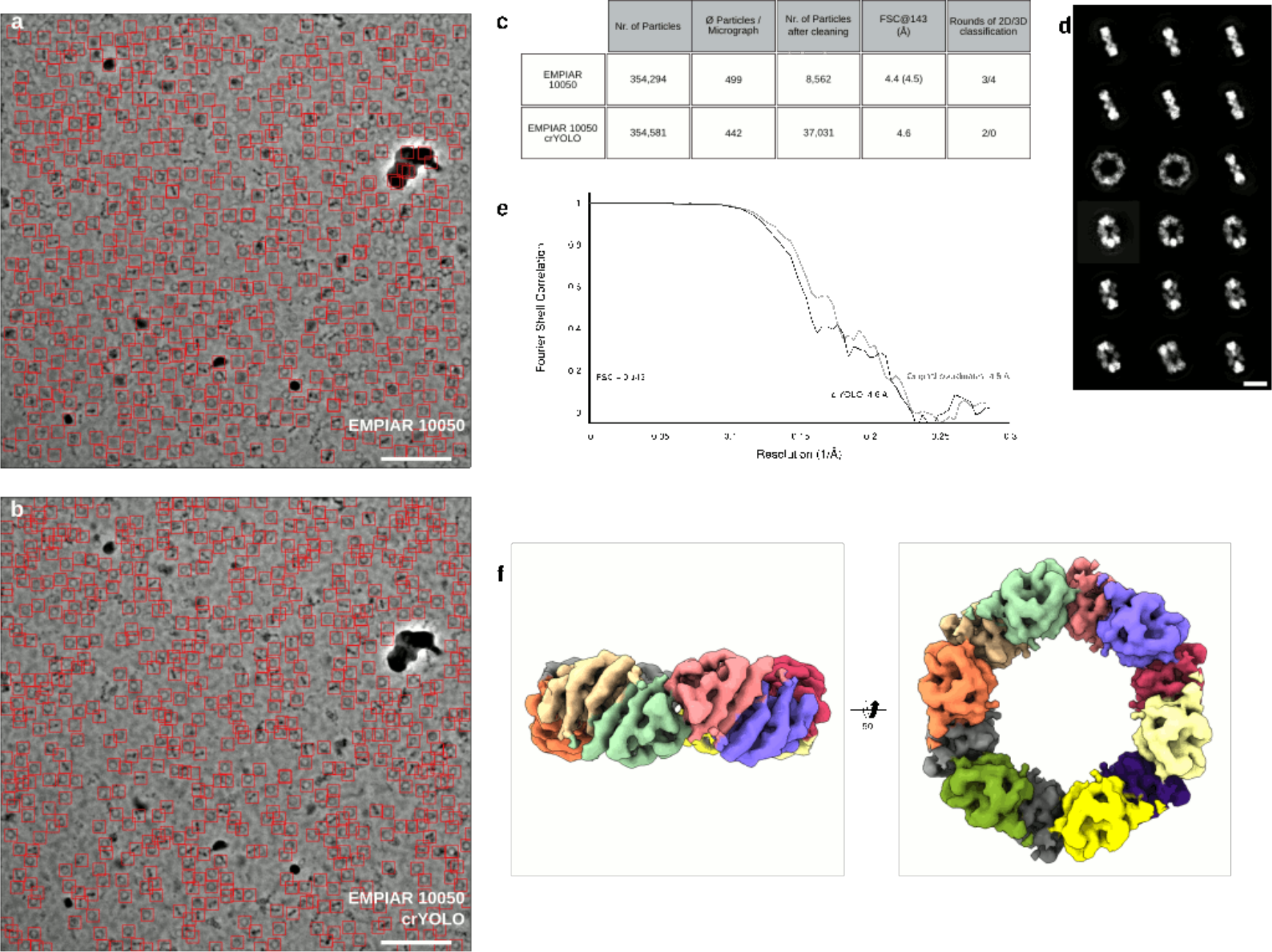
Selection of Prx3 particles and structural analysis. **a, b)** Particles selected on a representative micrograph (micrograph number 19.22.14) of the EMPIAR-10050 dataset using either **(a)** crYOLO or **(b)** EMAN2. Scale bar, 100 nm. **c)** Summary of particle selection and structural analysis. The resolution in parentheses is the result obtained after a 3-D refinement performed in SPHIRE, using the final 8,562 particles of the original dataset. **d)** Representative 2-D class averages obtained from 2 rounds of classification using the crYOLO-selected particles and ISAC. Scale bar, 10 nm. Well-centered examples for all views showing high-resolution details can be readily obtained from the data. **e)** Fourier shell correlation plots for the final 3-D reconstruction (black) using the crYOLO-selected particles or the 8,562 particles from the original dataset (gray). The average resolution of our 3-D reconstruction is ~ 4.6 Å while that one from the originally used particles is ~ 4.5 Å. **f)** Top and side views of the 3-D reconstruction obtained with crYOLO/SPHIRE. For clarity, all subunits are colored differently in the reconstruction.

The small differences in the final resolution of the reconstructions can be attributed to the fact that all reconstructions reach resolutions near the theoretical resolution limit; at a pixel size of 1.74 Å, 4.4 Å represents approximately 0.8 times Nyquist. Consistent with this observation, the FSC curves of the reconstructions from both sets of particles are practically superimposable (Figure 5e). Interestingly, a large fraction of the particles was discarded during refinement, both by the original authors – 97.6% – and by us – 87.7% kept. In our case, this is not due to the quality of the particle selection. Indeed, 310,123 of our picks were included into the stably aligned images after ISAC. Instead, we deleted most of the particles that were found in the preferred top orientation, which represents roughly half of the dataset. The remaining discarded images correspond to correct picks in regions where the protein forms clusters. In those cases, while the particles were correctly identified by crYOLO, they were not used for single particle reconstruction.

In order to evaluate the SNR dependency on the picking quality of crYOLO, we used the simulated data and trained a model for each noise level set, using the 15 micrographs for training and 4 for validation for each set. Finally, we calculated the AUC for each of the sets (Figure 6a). Up to a noise level of 6, the AUC value is greater than 0.8 and even for a noise level of 8 the AUC values greater than 0.6. In images with a noise level of 8 or greater, the particles are visually barely distinguishable from the background (Figure 6b). This demonstrates the strength of crYOLO for selecting particles of near-to-focus cryo-EM datasets that naturally have a low contrast.

**Figure 6.**
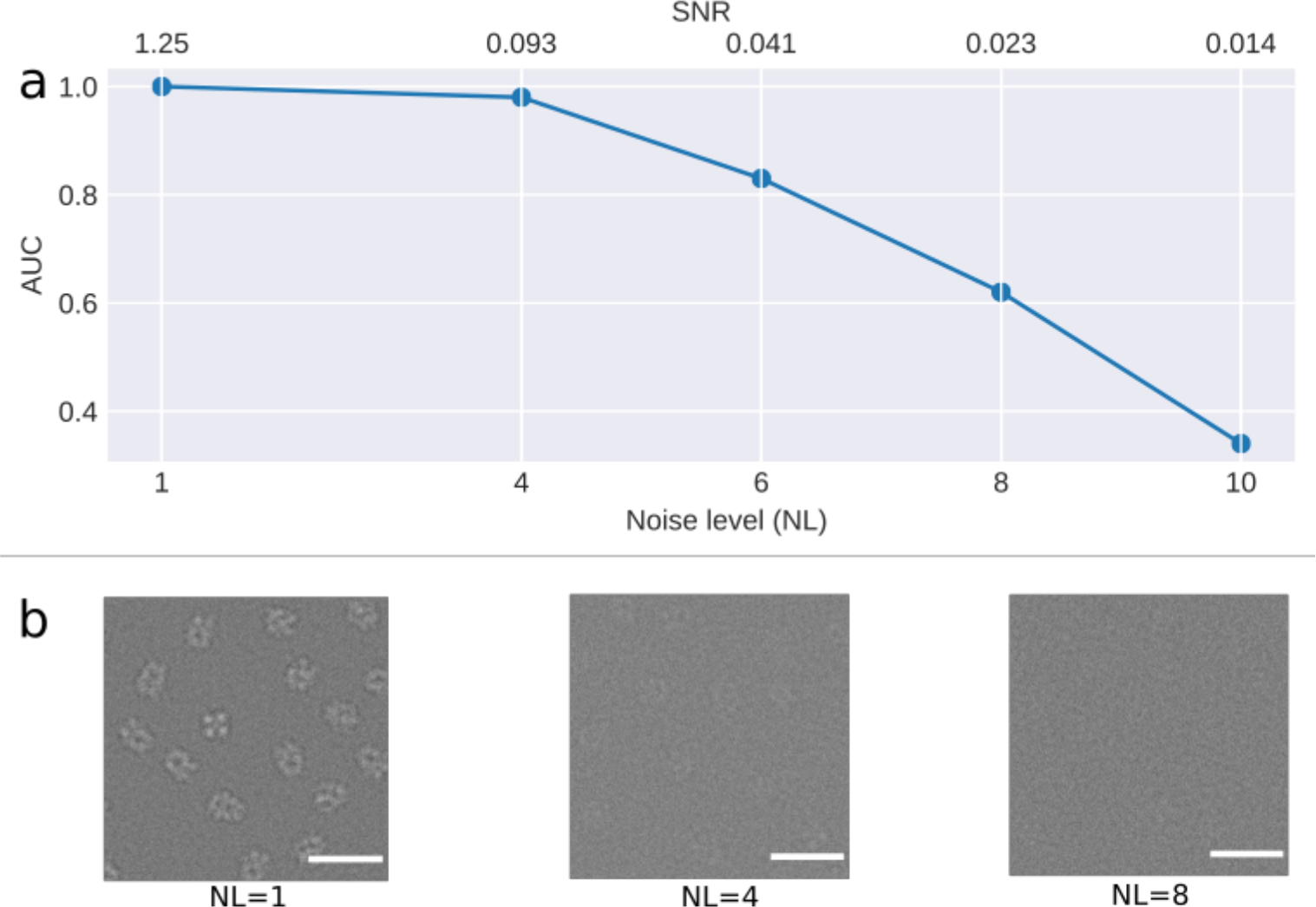
SNR dependence of crYOLO. **(a)** Noise level dependency of crYOLO picking simulated TRPC4 particles (EMD-4339) measured by the area under the precision-recall curve (AUC). The AUC stays above 0.8 up to a noise level of 6 (SNR 0.041). **(b)** Example micrographs for the noise levels of 1, 4 and 8.

To determine the influence of the size of the training dataset on the selection quality, we trained crYOLO on the manually picked KLH dataset and on subsets of different size. We decreased the number of micrographs in each subset gradually and evaluated the results by calculating the precision and recall scores. The selection performance was excellent, for both the high defocus and the low defocus micrographs (Figure 7a). To demonstrate the discrimination power, we trained a model to pick only the side views of KLH. The performance was comparable with picking all views (Figure 7b). Even if the model was trained with as few as 40 KLH particles from one micrograph pair, crYOLO still demonstrated a high AUC value of 0.9 (Figure 7c,d). All trained models skipped MAVS filaments, stacked KLHs and broken particles. This demonstrates that crYOLO can be easily and accurately trained with only few particles.

**Figure 7.**
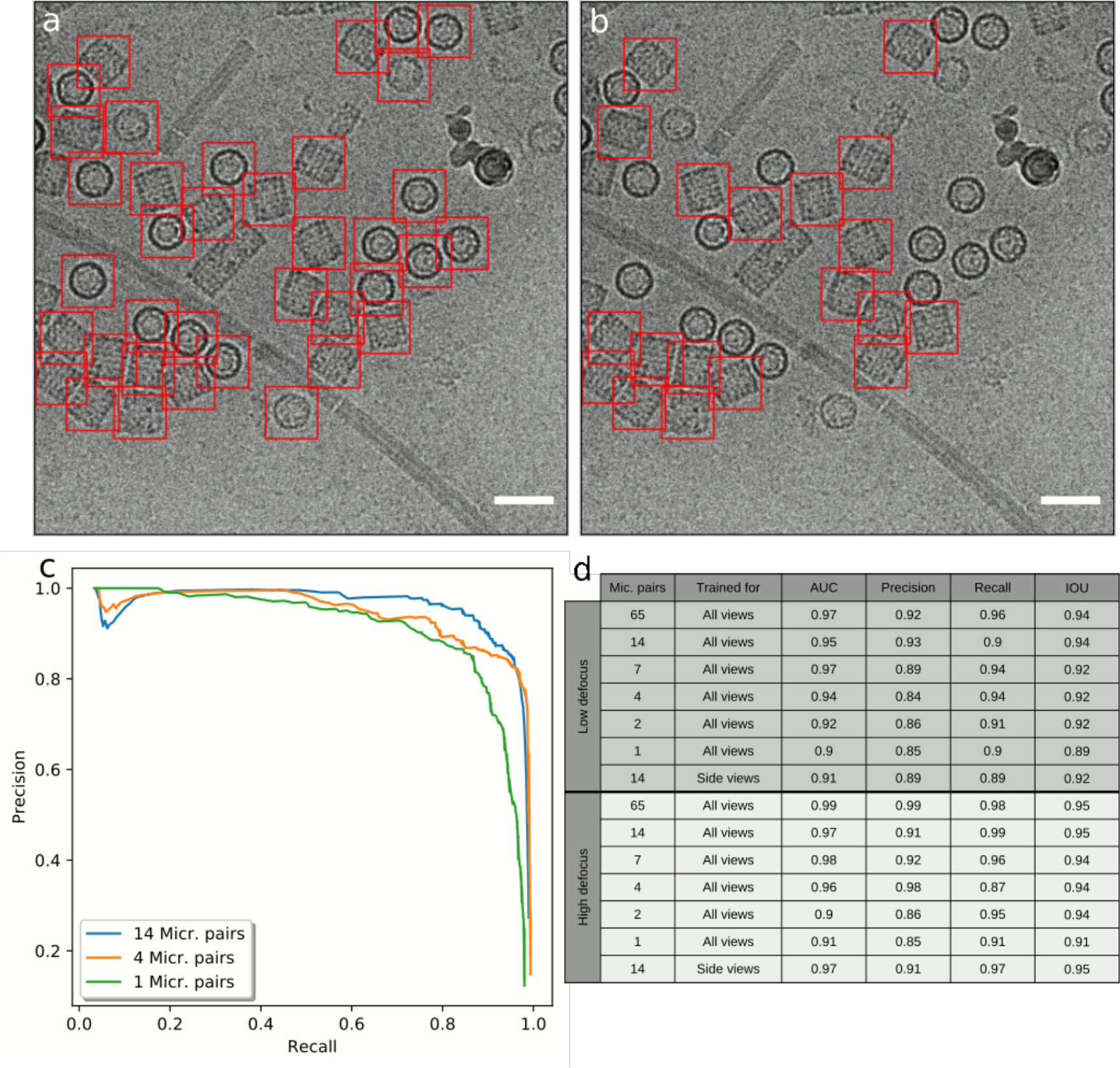
Training of crYOLO on KLH. **a)** One example of a particle picking result by crYOLO trained for all views with 14 micrographs of the full KLH dataset and (**b)** trained only for side views. Scale bar, 70 nm. **c)** Precision-recall curves for the low defocus micrographs of the KLH dataset using several training set sizes. The curves were estimated based on 17 randomly selected test micrographs out of the full dataset. The AUC values are 0.97 (blue), 0.94 (orange), 0.9 (green). **d)** Evaluation table for different subsets of the KLH micrographs. Additionally, a subset with 14 micrographs was trained to pick only the side views. The IOU and the AUC value of all experiments is between 0.89 and 0.97.

### Computational efficiency

We used a desktop computer equipped with a NVIDIA GeForce GTX 1080 graphics card with 8 GB memory and an Intel Core i7 6900K CPU to train crYOLO and select the particles using the GPU. The time needed for training was 5 to 6.5 minutes for each dataset (Figure 8). The selection process was very fast for all three datasets. The selection of the Prx3, TcdA1 and NOMPC datasets reached a speed of ~4.6, ~5.0 and ~4.2 micrographs per second, respectively. However, if no GPU is available, a trained model can also be applied on a common CPU. The picking speed on our multi-core CPU is one second per micrograph.

**Figure 8.**
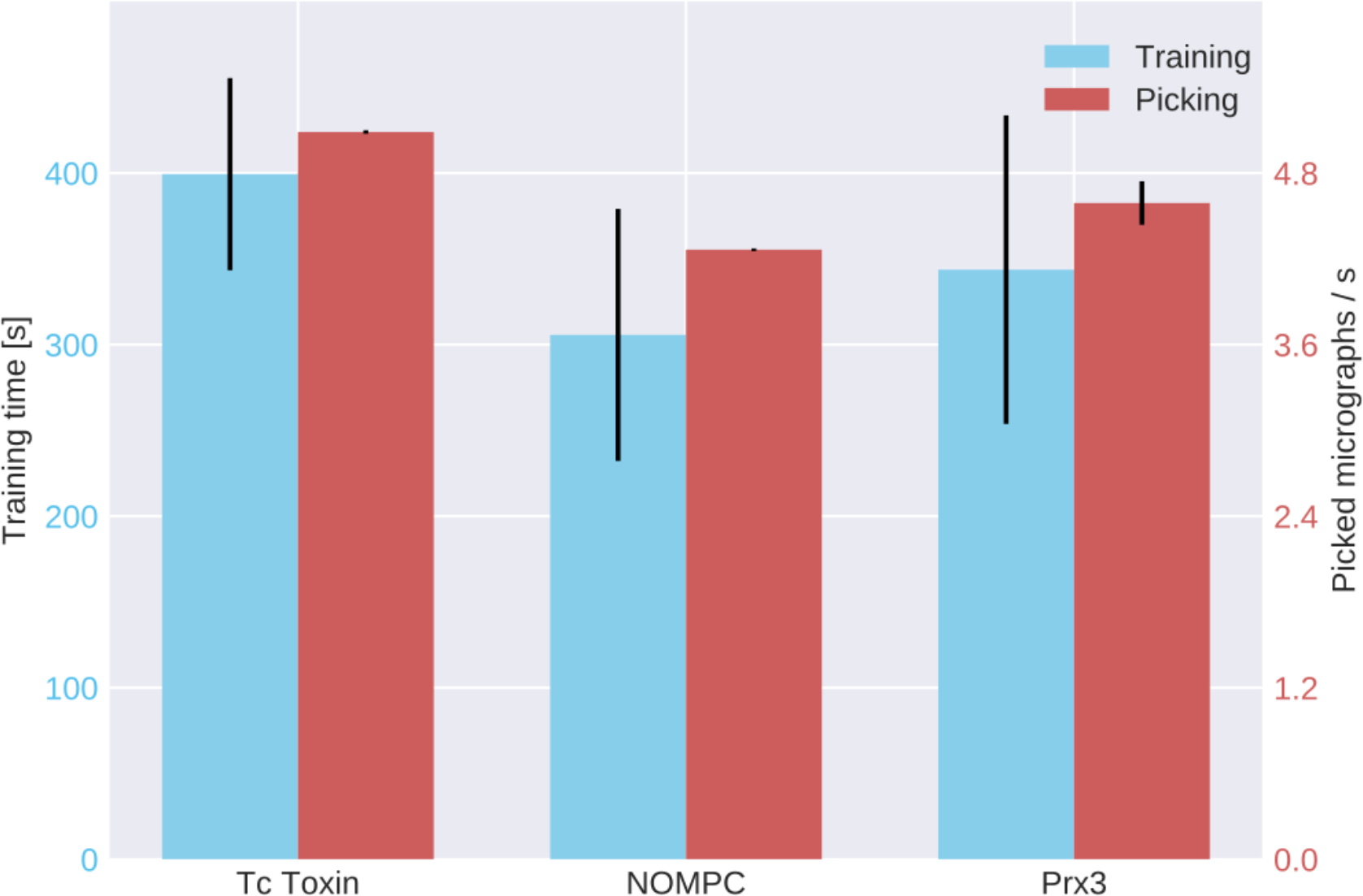
Computational efficiency statistics of training and particle selection. The crYOLO training time for TcdA1, NOMPC and Prx3 was 400, 300 and 343 seconds, respectively (blue bars). The red bars depict the number of processed micrographs per second during particle selection. Error bars are standard deviation, measured by training and applying crYOLO three times. All datasets were picked in less than a quarter of a second per image. For TcdA1, crYOLO needed 0.19 seconds per micrograph, for NOMPC 0.23 and 0.21 seconds for Prx3.

In comparison to other deep learning particle selection tools, this is quite fast. For example, the software DeepPicker^8^ reports 1.5 minutes per micrograph on a GPU and the software DeepEM^29^ reports 13 minutes on a CPU and 40 seconds on a GPU.

### Generalization to unseen datasets

Ideally, crYOLO would specifically recognize and select particles that it has not seen before. In order to reach this level of generalization, we trained the network with a combination of 840 micrographs from 45 datasets containing 26 manually picked datasets, 9 simulated datasets and 10 particle-free datasets consisting only of contamination. The molecular weight of the complexes from the respective datasets ranged from 64 kDa to 1.1 MDa.

Using this generalized crYOLO network, we automatically selected particles of RNA polymerase^30^ (EMPIAR 10190) and glutamate dehydrogenase (EMPIAR 10217). Although crYOLO had not been trained on these particles, it specifically selected particles while avoiding contamination (Figure 9a,b).

**Figure 9.**
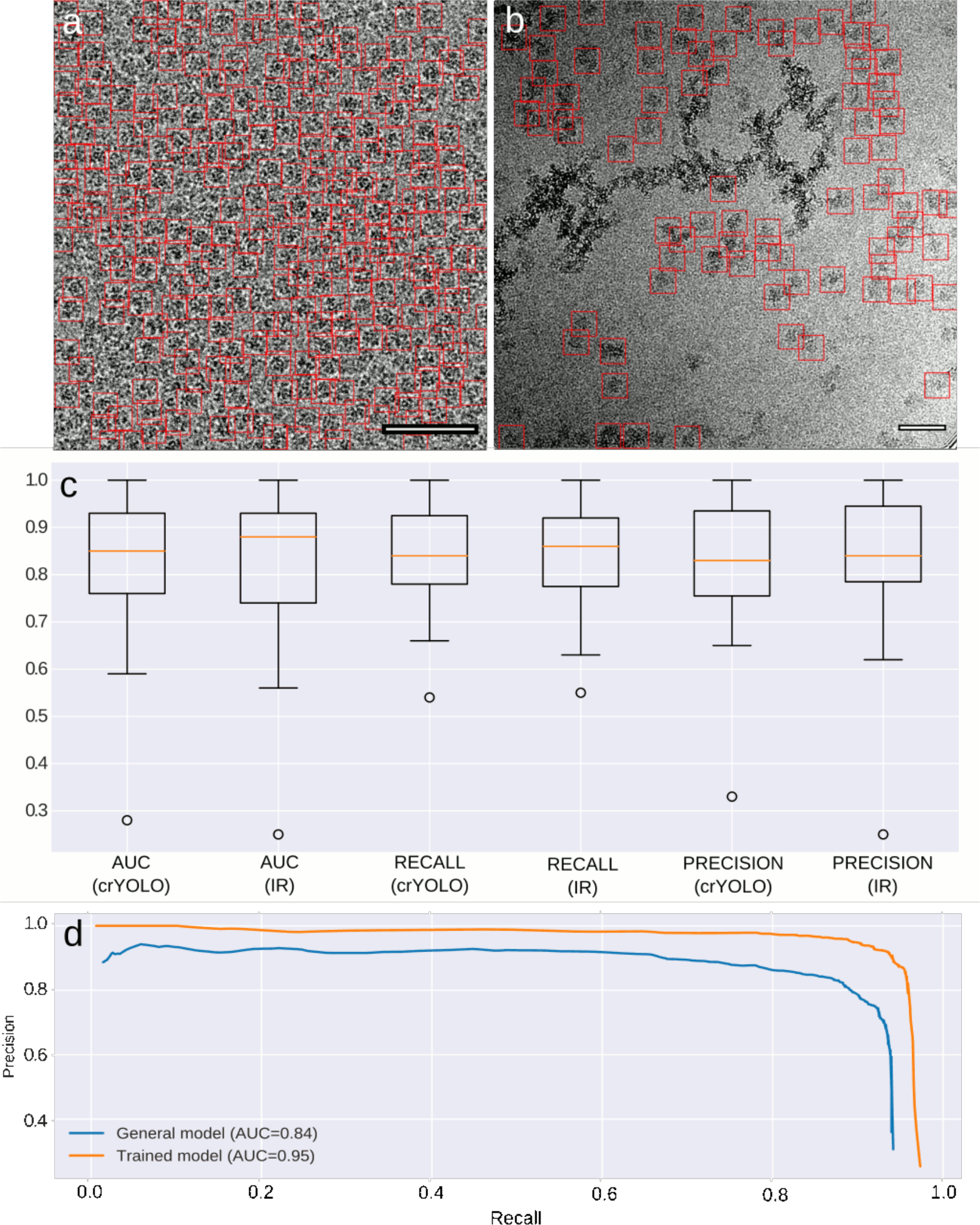
Generalized crYOLO network. **a, b)** Particles selected on a representative micrograph of glutamate dehydrogenase (EMPIAR 10127) and RNA polymerase (EMPIAR 10190). None of the datasets was included in the set used for training the generalized crYOLO network. Scale bars, 50 nm. **c)** AUC, recall and precision of the datasets included into the general model evaluated for the crYOLO network architecture and the Inception-ResNet (IR) architecture. The box shows the lower and upper quartiles with a line as median. The whiskers represent the range of the data, while the points are outliers. **d)** Precision-recall curves for TcdA1 picked with either a network directly trained on the TcdA1 dataset (orange) and the general model but not on TcdA1 (blue).

In the recent years, several new network architectures have been published. These networks have a higher capacity and allow the training of more than 100 layers. One of these networks is the Inception-ResNet^31^ (IR) which achieved high accuracy at the ImageNet competition^32^. To compare this network with the crYOLO network, we replaced the feature extraction part of crYOLO with the Inception-ResNet architecture and trained it on the complete combination of datasets. The IR architecture achieved the same performance as the crYOLO architecture (Figure 9c). As illustrated here, the median AUC/recall/precision is approximately the same. Thus, what we conclude is that, given our current training set, increasing the capacity of the network does not result in improved performance. Bigger networks have the advantage that they have even a higher capacity, i.e. can be trained to arbitrary depth. However, they tend to overfit more easily and the IR architecture comes with a higher computational cost. In addition, it was shown recently that many layers in modern ResNets do not learn anything^33^ and can be safely removed without damaging the performance of the network. This means that a network with the same prediction quality but lower number of weights, such as the one we used for crYOLO, is actually an advantage in a practical setup with limited training data. However, it is noteworthy to highlight that crYOLO offers the possibility of the integration of new architectures in a straight-forward manner, which will contribute to its sustainability.

To assess the quality of the generalized crYOLO network, we compared its performance selecting TcdA1 with that of a network that was directly trained on this protein. As expected, the AUC and localization accuracy are better for the directly trained network (Figure 9d). However, the generalized crYOLO network selected a similar number of particles (Figure 3d); the AUC of 0.84 (Figure 9d) and the IOU of 0.79 show that its performance is of sufficient quality to select a good set of particles. Indeed, the TcdA1 particles selected by the generalized crYOLO network, resulted in a reconstruction of similar resolution and quality as the particles from the TcdA1-trained crYOLO (Figure 3f). We expect that increasing the training set with a higher number of proteins and complexes will further expand the capabilities of crYOLO.

## Discussion

In this work, we present crYOLO, a novel fully-automated particle picking procedure for cryo-EM. CrYOLO employs a state-of-the-art deep learning object detection system which contains a 21-layer convolutional neural network. The excellent performance of crYOLO on several recent direct detector datasets, reflects the efficiency of the program to detect good particles with an accuracy comparable to manual particle selection at high speed. CrYOLO’s speed and efficiency underlines its potential to become a crucial component in a streamlined automated workflow for single particle analysis and thus eliminates one of the remaining bottlenecks.

To close the other remaining gaps in the automated workflow between the electron microscope and data processing, such as evaluation, drift and CTF correction, file conversion and transfer, our lab has developed an “on-the-fly” processing pipeline called TranSPHIRE, which is documented and freely available on www.sphire.mpg.de (manuscript in preparation). The TranSPHIRE pipeline includes crYOLO.

The crYOLO package is available as a standalone program but it is easy to be integrated in existing workflows. In the near future, it will be fully integrated into the SPHIRE image processing workflow and into the Scipion framework^34^. In addition to the command line interface, crYOLO provides an easy-to-use graphical tool for simple and direct generation of training data as well as visual evaluation of picking results (Figure 2). CrYOLO outputs the coordinate files in EMAN box-file and RELION STAR-file format which can easily be imported to all available software packages for single particle analysis. CrYOLO picks micrographs rapidly on a standard GPU. If a trained network is available, particle picking can also be performed on multi-core CPUs. However, this decreases the speed of selection to 1-3 seconds per micrograph. This is still faster than other particle pickers on a GPU. Some of these differences can be attributed to the exact model of CPUs and GPUs used. However, the use of the YOLO approach clearly plays a strong role in the particle picking time. Moreover, this approach is not only faster but has the capability to learn the context of the particle such as to avoid picking inside or in the vicinity of contamination. This feature is particularly important for specimens that pose a challenge due to the high amount of contamination present. In addition, what further contributes to the overall value of our crYOLO package includes the low number of micrographs that are required during the training process, along with eliminating the need for the user to pick negative examples.

Our evaluations demonstrate that the accuracy of crYOLO yields a set of particles with a remarkably low number of false positives. In addition, the IOU values of the selected particles provide evidence for the excellent centering performance of crYOLO. These are both strengths that contribute to improving the quality of the input data set for image processing, and ultimately improve the final reconstruction in many cases. At the same time, it reduces the number of subsequent processing steps, such as computationally intensive 3-D classifications, and thereby shortens the overall processing time.

In comparison to template-based particle picking approaches, crYOLO is not prone to template bias and therefore the danger of picking “Einstein from noise”^35^ is reduced. However, the user should select the particles for the training data without bias. Otherwise the applied supervised learning method will create a biased model. For example if a view of the particle is completely missing in the training data, crYOLO might learn to skip this view. Therefore the micrographs in the training dataset should be representative of the full dataset with respect to defocus, types of contamination etc.

Importantly, crYOLO can be trained to select previously unseen datasets. With the use of a training set containing 10-20 micrographs from 40 datasets, we have obtained a powerful general network. Adding more training datasets from different projects that were manually picked by experienced users will improve the performance of crYOLO even further. In crYOLO adding more training data is especially easy, because in contrast to the sliding-window approaches, training crYOLO only requires particles to be labeled and not the background or any contamination. Therefore, previously processed datasets can be directly used to train crYOLO without manually picking background and contamination afterwards. In principle, it is also feasible to train generalized networks that are specialized on certain types of particles, such as viruses, elongated or very small particles to improve the performance of crYOLO. At the moment, the development of convolutional neural networks is moving fast and new CNN networks which outperform earlier networks are released frequently. The flexible software architecture of crYOLO allows an easy incorporation of new CNNs and therefore a straightforward adaptation to new developments, if necessary.

Taken together, crYOLO enables rapid fully-automated “on-the-fly” particle selection with comparable precision as manual picking, without bias from reference templates. Furthermore, with the use of the generalized model presented here, our particle picker, crYOLO, can be used without a template or human intervention to select particles on most single particle cryo-EM projects within a very short period of time.

## Acknowledgments

We thank Yifan Cheng and Radostin Danev for kindly providing us the original coordinates of the NOMPC and Prx3 particles, respectively. This work was supported by the Max Planck Society (S.R.) and the European Council under the European Union’s Seventh Framework Programme (FP7/2007–2013) (Grant 615984 to S.R.).

## Contributions

S.R. designed the project, T.W. conceived the approach and wrote the software. T.W., C.G., M.S., T.M, optimized the approach, T.W., C.G., F.M. evaluated the performance, T.W., C.A, A.A., P.H., O.S, T.R, T.S, D.P, D.Q, S.T., B.S, E.S., and P.L picked datasets for the general model, T.W. and S.R. wrote the manuscript with input from all authors. All authors discussed the results and the manuscript.

## Data availability

The training datasets for this study are available from the corresponding author upon reasonable request. CrYOLO – along with a detailed practical manual - is available for download under http://sphire.mpg.de

## Online Methods

### CrYOLO architecture and training

CrYOLO trains a deep convolutional neural network (CNN) for automated particle selection. A typical CNN consists of multiple convolutional and pooling layers, and is characterized by the depth of the network, which is the number of convolutional layers^36^. The input of a CNN is data with a grid-like topology, most often images which may have multiple channels (e.g. color). A convolutional layer consists of a fixed number of filters, and each filter consists of multiple convolutional kernels with a fixed size (typical 1×1, 3×3 or 5×5). The number of convolutional kernels per filter is equal to the number of channels in the input data of the layer. During forward propagation, each filter in a convolutional layer creates a feature map which is a channel in the final output of the layer. Each kernel is convolved with the corresponding input channel. The results are summed up along the channels, resulting in the first channel of the output. The same procedure is applied for the second filter which results in the second channel and so on. After all channels are calculated, the so-called batch normalization^37^ is applied. Batch normalization normalizes the feature map and leads to faster training. Furthermore, it has some regularization power which makes further regularization often unnecessary. Finally, a non-linear activation function is applied element-wise to the normalized feature map which results in the final feature map.

In typical CNNs, max pooling layers are inserted between some of the convolutional layers. Max pooling layers divide the input image into equal sized tiles (e.g. 2×2) which are then used to calculate a condensed feature map. Therefore, for each tile a cell is created, the maximum for the tile is computed and inserted into the cell. This leads to a reduced dimensionality of the feature map, and makes the network more memory-efficient and robust against small perturbations in the pixel values.

The network architecture used in crYOLO is summarized in Table 1. The feature extraction part consists of 21 convolutional layers and each convolutional layer consists either of multiple 3×3 filters or 1×1 filters. In addition, it contains a passthrough layer between layer 13 and 21, which enables the network to use low level features during prediction. All convolutional layers use padding so that they do not reduce the size of their input. We used the leaky rectified linear unit^38^ (LRELU) as activation function. LRELU simply returns the element itself if the element is positive and the element multiplied by a fixed constant α (here α=0.1), if it is negative.

**Table 1.** YOLO network architecture. The CNN consists of 21 convolutional and five max-pooling layers for feature extraction. For detection, a single convolutional layer is used with a dropout layer in front to reduce overfitting. The dropout layer is only used during training, not during prediction.

| Layer | Type | Filters | Size |
| --- | --- | --- | --- |
| Feature Extraction |  |  |  |
| 1 | Convolutional | 32 | 3x3 |
|  | Max-Pooling |  |  |
| 2 | Convolutional | 64 | 3x3 |
|  | Max-Pooling |  |  |
| 3 | Convolutional | 128 | 3x3 |
| 4 | Convolutional | 64 | 1x1 |
| 5 | Convolutional | 128 | 3x3 |
|  | Max-Pooling |  |  |
| 6 | Convolutional | 256 | 3x3 |
| 7 | Convolutional | 128 | 1x1 |
| 8 | Convolutional | 256 | 3x3 |
|  | Max-Pooling |  |  |
| 9 | Convolutional | 512 | 3x3 |
| 10 | Convolutional | 256 | 1x1 |
| 11 | Convolutional | 512 | 3x3 |
| 12 | Convolutional | 256 | 1x1 |
| 13 | Convolutional | 512 | 3x3 |
|  | Max-Pooling |  |  |
| 14 | Convolutional | 1024 | 3x3 |
| 15 | Convolutional | 512 | 1x1 |
| 16 | Convolutional | 1024 | 3x3 |
| 17 | Convolutional | 512 | 1x1 |
| 18 | Convolutional | 1024 | 3x3 |
| 19 | Convolutional | 1024 | 3x3 |
| 20 | Convolutional | 1024 | 3x3 |
| 21 | Convolutional | 1024 | 3x3 |
| Detection |  |  |  |
|  | Dropout (0.2) |  |  |
| 1 | Convolutional | 6 | 1x1 |

Five max pooling layers downsample the image by a factor of 32. During training, a dropout unit in the detection part sets 20 percent of the entries in the feature map after layer 21 to zero. This regularizes the network and reduces overfitting. Finally, a convolution layer with six 1×1 filters performs the actual detection. With the YOLO^16^ approach, each cell in the final feature map is used (i) to classify if the center of a particle box is inside this grid cell and (ii) if this is the case, it predicts the exact position of the box center relative to the cell, as well as the width and height of the box.

The network was trained using backpropagation with the stochastic optimization procedure ADAM^39^. Backpropagation applies the chain rule to compute the gradient values in every layer. The gradient determines how the kernel elements in each convolutional layer should be updated to get a lower loss. The optimizer determines how the gradient of the loss is used to update the network parameters. For YOLO, the loss function to be minimized is given by:

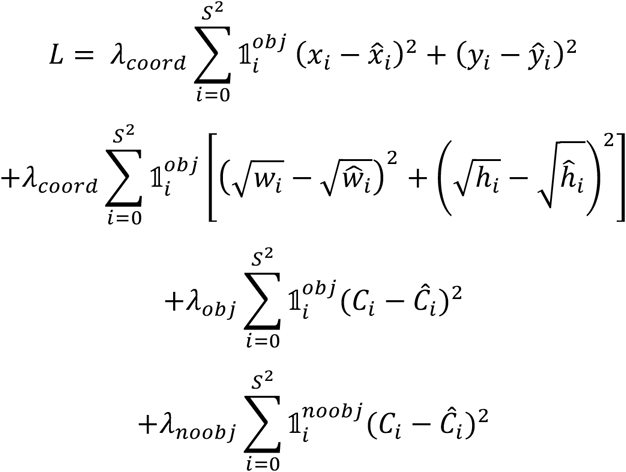

where 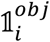 is 1 if the center of a particle is in cell *i* and 0 otherwise, λ_*coord*_, λ_*obj*_ and λ_*noobj*_ constant weights, (x_*i*_,y_*i*_) the centrum coordinates of the training boxes and 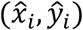 the predicted coordinates. The width and height of the box with their estimates are given by (w_*i*_,h_*i*_) and 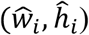 respectively. The confidence that a cell contains a particle is *C*_*i*_.

The first term of the loss function penalizes bad localization of particle boxes. The second term penalizes inaccurate estimates for the width and height of the boxes. The third term attempts to increase the confidence for cells with a particle inside. The last term decreases the confidence of those cells containing no particle center. The loss function slightly differs from the one used in Redmon et al.^12^, since we only have a single class to predict and also only one reference box (anchor box in Redmon et al.^12^).

### Data Augmentation

During training, each image is augmented before passing it through the network. This means that it is slightly altered by random selection methods instead of passing the original image through the network. Each image is passed multiple times through the network, randomly modified in different ways. This helps the network to reduce overfitting and also the amount of training data needed. The applied methods are:

- Gaussian blurring: a random standard deviation between 0 and 3 is selected and then a corresponding filter mask is created. This mask is then convolved with the input image.
- Average blurring: a random mask size between 2 and 7 is chosen. This mask is shifted over the image. At each position, the central element is replaced with the mean values of its neighbors.
- Flip: the image is mirrored along the horizontal and vertical axes.
- Noise: Gaussian noise with a randomly selected standard deviation proportional to the image standard deviation is added to the image.
- Dropout: randomly replaces 1 to 10% of the image pixels with the image mean value.
- Contrast normalization: the contrast is changed by subtracting the median pixel value from each pixel, multiplying them by a random constant and finally adding the median value again.

